# Pair-wise Interactions in Gene Expression Determine a Hierarchical Transcription Profile of the Human Brain

**DOI:** 10.1101/2020.05.15.098962

**Authors:** Jiaojiao Hua, Zhengyi Yang, Tianzi Jiang, Shan Yu

## Abstract

Orchestrated expressions of tens of thousands of genes give rise to the complexity of the brain. However, it is unclear what is the structure governing these myriads of gene-gene interactions. By analyzing the transcription data obtained from more than 3000 sites in human brains, we found that pair-wise interactions between genes are sufficient to accurately predict both the transcription pattern of the whole genome for individual brain areas and the transcription profile of the entire brain, suggesting a surprisingly simple interaction structure of the transcriptome itself. We further revealed that the strength of gene-gene interaction observed empirically allows the nearly maximal number of transcriptionally similar clusters of areas to form, which may account for the functional and structural richness of the brain.

**One Sentence Summary:** Pairwise interactions among genes shape the overall structure of human brain transcriptome.

## Introduction

The human brain contains hundreds of anatomically and functionally diverse areas, each of which is composed of billions of heterogeneous cells. In addition, there are complex anatomical connections as well as functional interactions that coordinate the entire brain network to perform various functions. Such a complex, hierarchical structure is highly stereotyped between individuals, suggesting a common underlying transcriptional regulation. Indeed, the study of human brain transcriptome has provided important insights into how orchestrated expression of numerous genes control the structure, function, development, and diseases of the brain [1].

Specifically, it has been reported that transcriptional profiles can strongly reflect the spatial topography of the neocortex, manifested as higher similarity of the transcriptional profiles for spatially close regions [2] or adjacent cortical layers [3]. Also, the consistency of transcriptional pattern was found to be correlated with resting state functional connectivity among different brain areas [4]–[7], especially with the connectivity extracted by the low-frequency fluctuations in the BOLD signals [5]. Moreover, gene expressions have been found to be related with functional difference of the brain either within the same or among various species [8]–[13]. Many specific expression patterns have been identified as critical controller that drive brain development [14]–[17]. In addition, it has been widely recognized that gene expression network plays a vital role in many psychiatric disorders such as autism, schizophrenia, bipolar disorder, Alzheimer’s disease, etc. [2], [18]–[24]

So far much attention has been focused on using the correlational approach to identify the role played by low-dimensional features of the transcriptome, e.g., the state(s) of specific genes or gene modules. However, it is known that the phenotype is determined by the complete pattern of gene expression [25], which cannot be fully revealed by studying the aggregated effects of individual genes or gene modules. The pattern of entire human brain transcriptome is extremely complex. At the microscopic level, the expression of tens of thousands of individual genes interacts with each other to determine the expression pattern of the genome [26]. At the macroscopic level, the expression patterns of genome among hundreds of distinct brain areas give rise to the transcription profile of the entire brain. At each level, there could be interactions involving pairs, triplets, quadruplets, etc. of genes or areas, leading to the complexity of 2^*N*^, where *N* is the number of genes or brain areas. Such high complexity limits the effectiveness of the traditional correlational approaches, impeding a comprehensive understanding that links gene expression to brain’s development, structure, function, and diseases.

Through analyzing the human brain transcriptome provided by the Allen Institute for Brain Science (AIBS) [2], here we found that pair-wise interactions between genes can accurately predict both the transcription pattern of the whole genome for individual brain areas, and the transcription profile of the entire brain, thus reducing the complexity of the human brain transcriptome dramatically, i.e. from 2^*N*^ to *N*^*2*^. It opens the possibility for further studies to comprehensively examine the function of genes by analyzing complete expression patterns.

Moreover, it enabled us to establish a quantitative framework to link individual gene expressions to the overall transcriptional clustering of the entire brain network. We found that the strength of gene-gene interaction observed empirically in the brain leads to nearly maximal number of clusters of areas, which may account for the functional and structural richness of the brain.

## Results

The dataset contains continuous (see Fig. 1A-B for examples) as well as binarized (Fig. 1C; 1 and 0 represent active and inactive expression states, respectively) transcriptional data of the human brain from six subjects (in total there were 3702 sample sites distributed across the whole brain). From this dataset, the expression levels of 16906 genes were obtained [27], allowing us to examine the interaction structure among individual genes. Further, to study the expression patterns across different brain areas, the samples were registered to 234 regions according to the Human Brainnetome Atlas [28] (Fig. 1A).

**Fig. 1.**
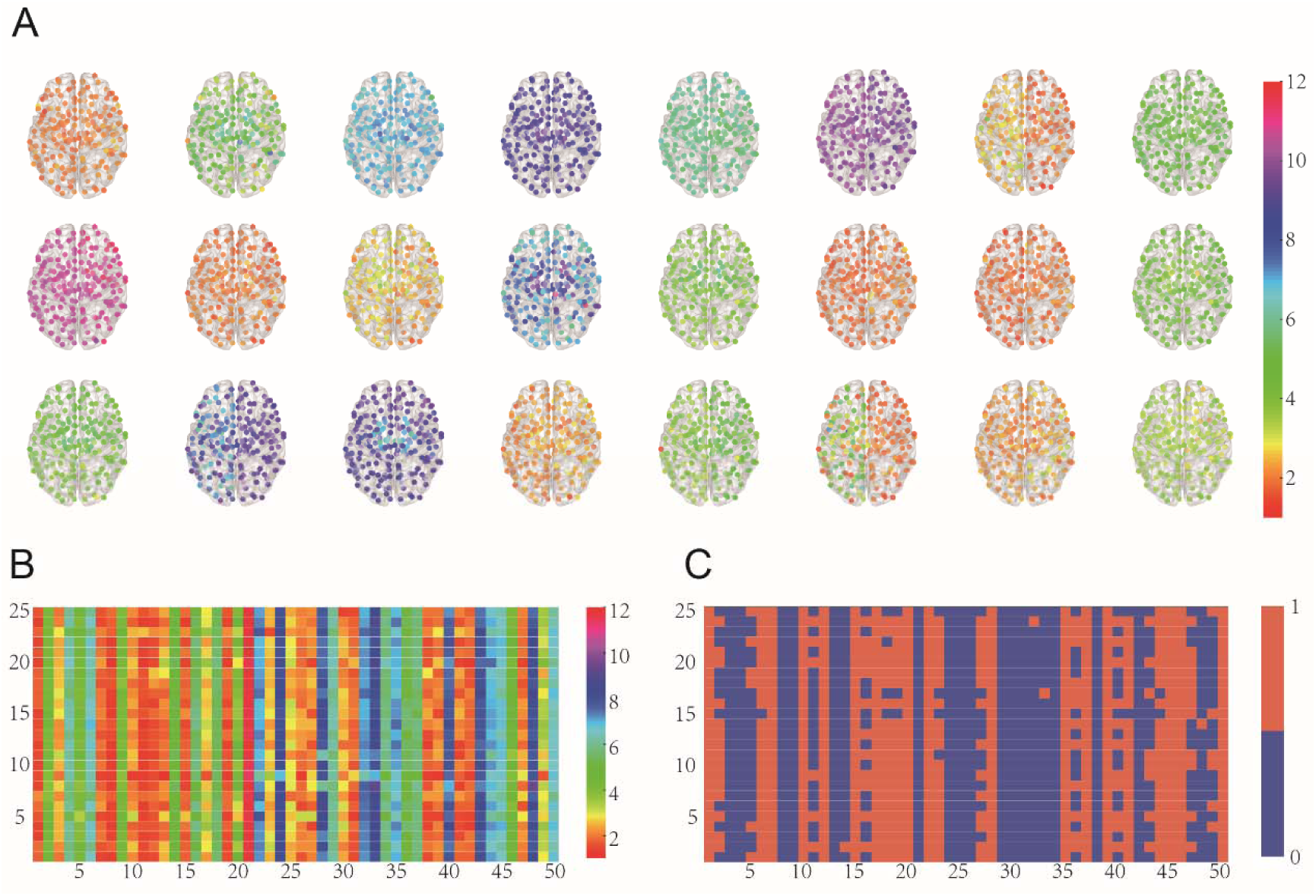
Examples of gene expression patterns. **(A)** Illustrations of how a single gene expresses differently across 234 regions. The expression patterns of 24 exemplary genes are shown, with each dot representing one area. The continuous gene expression level is color-coded. **(B-C)** Illustration of the continuous (B) and binarized (C) expression pattern of 50 genes across 25 regions. Both genes and regions were chosen randomly. Color bar represents the continuous or binarized gene expression value.

### Analysis on the small datasets

Here, we randomly chose small sub-dataset like a 6-gene and 6-area system and compared the possibility of the 10 genes’ or 10 areas’ expression patterns (“000000” ∼ “111111”) from the true data and the model prediction separately. The Independent Model [29] and the Dichotomized Gaussian Model (the DG model) [30], [31] were used here (See methods for details). The underlying rationale is the maximum entropy principle (Dewar, 2003; Jaynes, 1957; Lezon et al., 2006; Schneidman et al., 2006), i.e., the model that accurately predicts the expression patterns with the least information represents the true interaction structure among genes or areas.

#### The transcription pattern of 6 genes for individual brain areas

Firstly, at the gene level, we examined how the expression pattern of a population of genes emerges from the interactions among individual genes. To this end, We started from randomly choosing a population of *N* = 6 genes and examined different models’ performance in predicting the probability of expression patterns of 6 genes. We found that the prediction from the Independent Model, i.e., assuming expressions were independent among genes, was not as good as that from the DG model, i.e. incorporating the pairwise expression covariance (Fig. 2A). We repeated chose a population of 6 genes1000 times and found similar result in each case. Also, we obtained the distribution of JS divergence [32] of the possibility of expression patterns between the true data and DG model and between the data and the Independent Model, and found the Jensen-Shannon (JS) divergence between the data and DG model is much closer than that between the data and the Independent Model (Fig. 2B). These results suggested that the gene-gene interactions are important in determining the expression pattern of 6 genes. Next, we defined a feature, relative entropy drop (RED), to measure how much of the interaction structure of the 6-gene system can be captured with the DG model (See methods for details). The RED of the DG model was around 78.4% ±13%. Then, we chose other different *N* = (7, 8,…,18), and found that with the increase of *N*, the corresponding RED is increasing all the way around 85% (Fig. 2C). We also performed the process on all 2124 sample sites instead of mapping them to regions. With more samples, RED increased up to around 93% (Fig S8), indicating that the expression pattern of a population of genes is determined by pairwise interactions between these genes.

**Figure 2.**
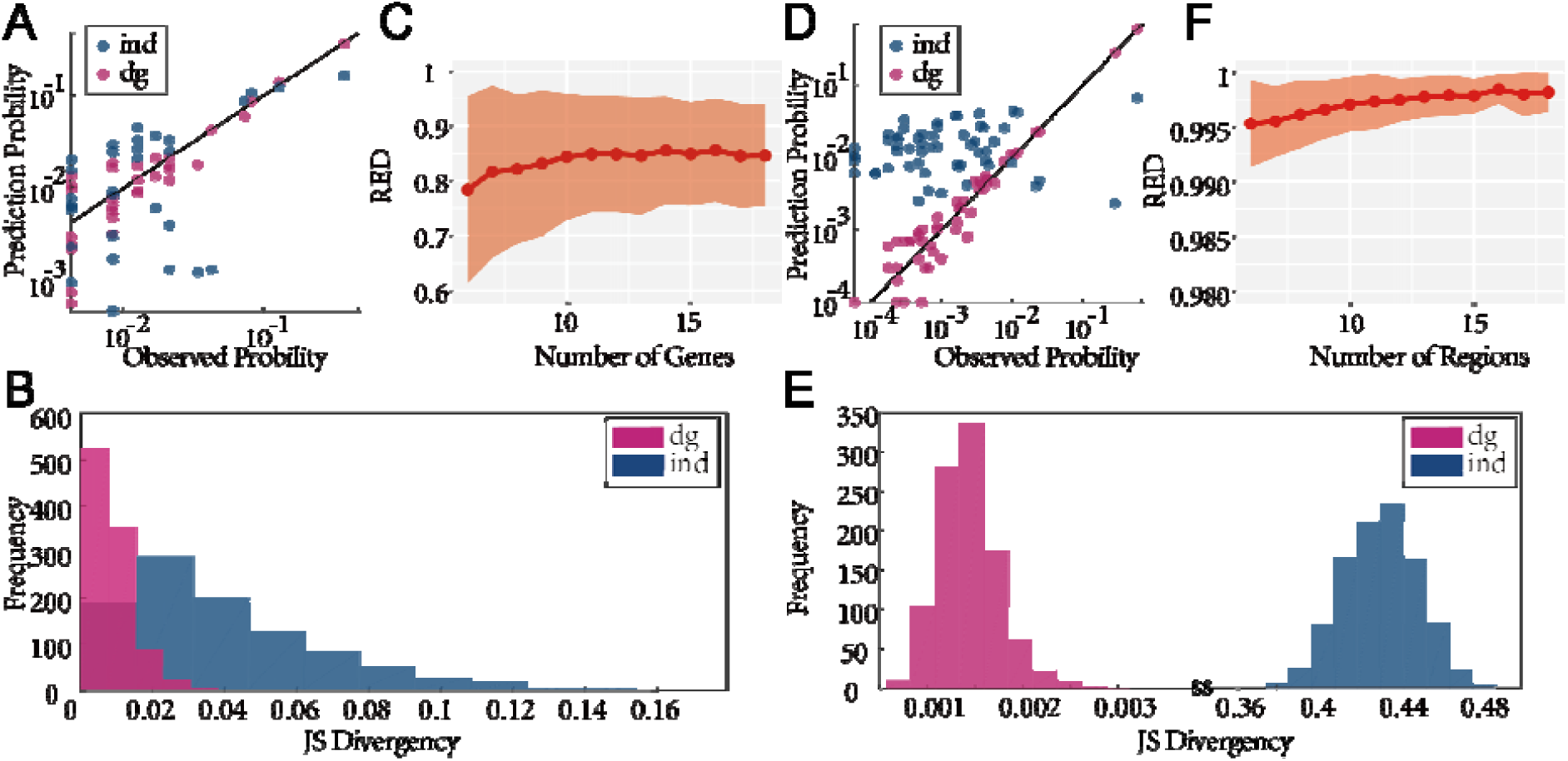
The expression pattern of 6-gene and 6-area system can be predicted by pair-wise interactions between genes and areas, respectively. **(A, D)** using random chosen 6 genes (A) or 6 areas (D), the possibility of each expression pattern observed is plotted against that predicted from the DG model (red dots) and IM (blue dots). Black line shows equality. **(B, E)** Distributions of JS divergences between the possibility of expression patterns observed and that predicted by DG model (red) and IM (blue); data is from 1000 groups of randomly chosen 6 genes (B) or 6 areas (E). **(C, F)** Randomly chose 6∼18 genes (C) or areas (F), 1000 times each. Number of genes are plotted against REDs. The line and shaded areas represent the mean ± variance, respectively.

#### The transcription profile of 6 areas

Next, at the area level, we studied how interactions of gene expression among different areas can give rise to the expression profile of a population of areas. Similar to the approach used above, we started from small dataset and randomly chose 6 areas. We found the probability of expression pattern of the 6 areas predicted by the DG model, adding the pair-wise interactions between areas, is much closer to the true data, comparing with the prediction from the Independent Model (Fig. 2D). As well, we found the distribution of JS divergence between DG model and the true distribution is much smaller in the 1000 times simulation (Fig. 2E). These results suggested that the area-area interactions are important in determining the expression pattern of 6 areas. In addition, the RED of the DG model is around 99.5% ± 0.39% (Fig 2F), suggesting that expression pattern of a population of areas is determined by pairwise interactions between these areas.

### Analysis on bigger datasets

To examine the robust of the result obtained from the small dataset, we used much bigger datasets, include 4074 genes and 234 areas. Here, we got rid of the genes which is active or inactive in most of the areas. Similarly, we still used the Independent Model and DG model and studied from two levels, the gene and area level respectively. At the gene level, we used two features of genomic expression pattern to evaluate the performance of different models: 1) the distribution of the proportion of active genes (PAG) in each brain area (Fig. 3A top); 2) the distribution of correlation coefficient of genome expression (CGE) between two areas (Fig. 3A bottom). At the area level, additional two features of genomic expression pattern of the brain were used: 1) the distribution of the proportion of brain areas in which a specific gene was active (PAA, proportion of active area), and 2) the distribution of correlation coefficient of expression of two genes across brain areas (CEA, correlation of gene expression) (see Fig. 3B for illustration of the PAA and CEA).

**Fig. 3.**
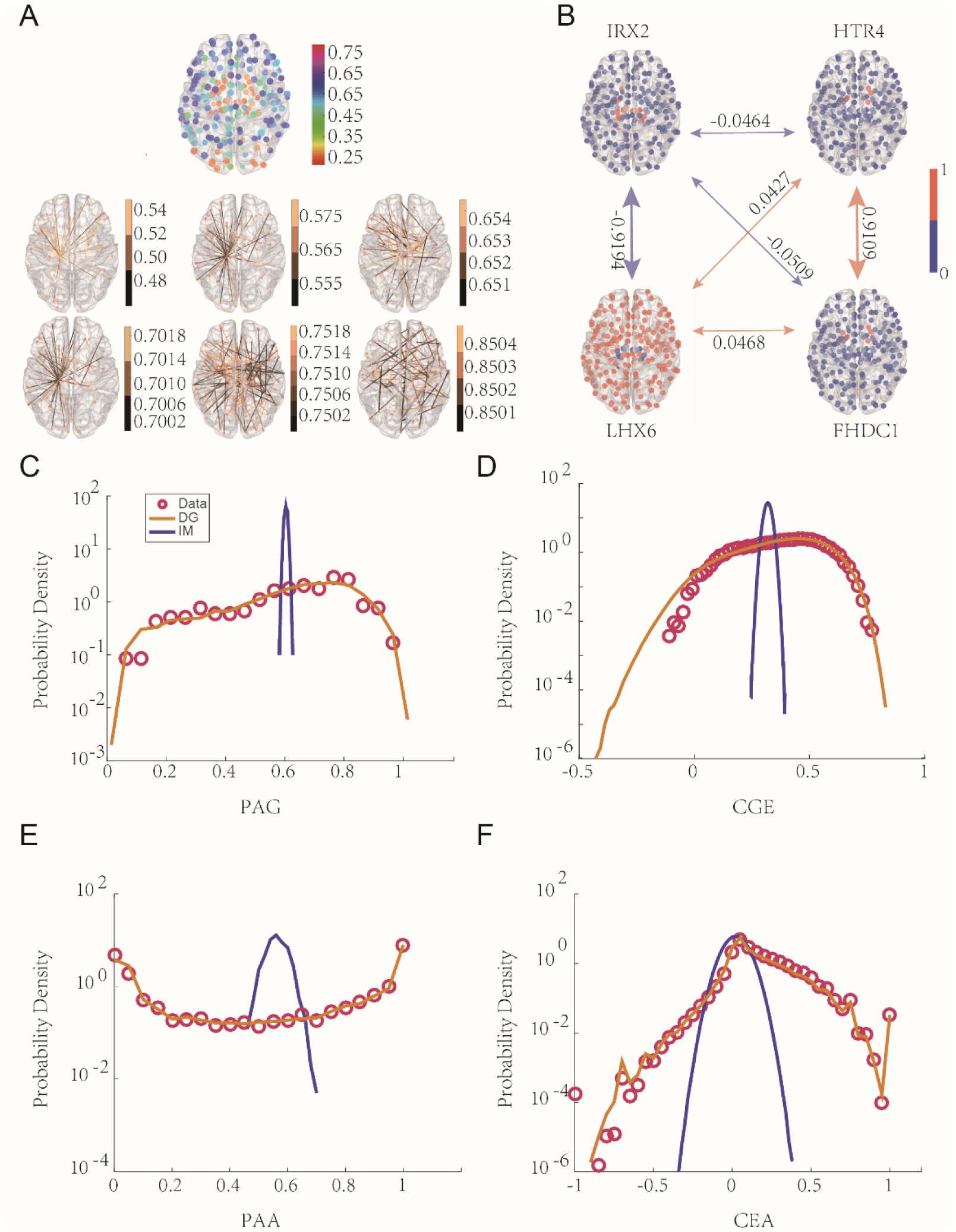
The expression pattern of individual brain areas and the whole brain can be accurately predicted by considering pair-wise interactions between genes and areas, respectively. **(A)** Top, an illustration of different gene expression profile of all brain areas. Each dot represents the proportion of active genes (PAG), which is color-coded. Bottom, illustrations of correlation of genome expression between areas (CGE). CGEs are represented by edges connecting corresponding areas, and the value is color-coded. For visual clarity, CGEs of different ranges are shown in separated panels. **(B)** Illustrations showing the correlation of single gene’s expression pattern. Each panel shows the proportion of brain areas where a specific gene was active (PAA). The binary expression level is color-coded. Edges connecting different panels (red, positive correlation, blue, negative correlation) show the correlation coefficient of expression patterns of two genes (CEA). **(C-F)** The empirically observed distributions of PAG, CGE, PAA, CBA (red circles) and predictions based on the IM (blue line) and the DG model (yellow line).

#### The transcription pattern of the whole genome for individual brain areas

At the gene level, we examined how the expression pattern of the whole genome emerges from the interactions among individual genes. We found that brain areas exhibited diverse levels of the overall gene expression, resulting in a unimodal but fairly broad distribution of the PAG (Fig. 3C). Meanwhile, genomic expression patterns between two areas tended to be correlated positively but distributed across a wide range (Fig. 3D). These high-level, quantitative measures reflect important features of genomic expression patterns. By examining different models’ performance in predicting these features, we found that only taking the probability of each gene to be active into account, i.e., assuming expressions were independent among genes (the Independent Model[29][29](Schneidman et al., 2006)), completely failed to predict the distributions of both the PAG and CGE. Indeed, by incorporating the pairwise expression covariance into the DG model, the distributions of the PAG and CGE could be accurately predicted (Figs. 3C, D, S2-3, S6), suggesting that the gene-gene interactions are important in determining the expression pattern of the genome. These results indicate that the expression pattern of the genome is determined by both the characteristics of individual genes, i.e., how likely a single gene is to be active, and the pairwise interactions between genes. This finding is surprising as tens of thousands of genes could have the triplet, quadruplet or even more complex interactions. However, it seems that all these higher-order interactions almost play no significant role in determining genome expression patterns.

### The transcription profile of the entire brain

At the area level, we studied how interactions of gene expression among different areas can give rise to the expression profile of the entire brain. In Fig. 3E, PAA showed a U shape distribution. A large proportion of genes expressed in a majority of areas, resulting in a concentrated region on the right side of the distribution. Not unexpectedly, among them, we identified abundant housekeeper genes [33]–[37]. Meanwhile, a similarly high proportion of genes resided on the left side of the distribution, indicating that many genes were inactive in most of the areas. In addition, the distribution of CEA showed a peak near 0 (Fig. 3F), demonstrating that the expression patterns between different gene were much less correlated compared to genome expression patterns between different areas. Similar to the results obtained at the gene level, we found that both the distribution of PAA and CEA can be predicted accurately by considering the observed covariance of the genome expression between areas (Fig. 3E, F, S4-6), suggesting a simple pair-wise areal interaction structure underlying the seemingly complex expression profile of the whole brain.

Consistent with the above analysis that the pair-wise structure can preserve the first and second order statistics when new elements are added or removed, we found that at both the gene and area levels, the distribution of the first and second order statistics are kept constant regardless of the number of genes/areas taken into account. Specifically, at the gene level, we randomly selected 500, 800, 1000 genes 5 times each, and obtained the distribution of PAA and CEA (Fig. 4A-B). At the area level, we randomly chose 100, 150, 250 regions 5 times each and obtained the distribution of PAG and CGE (Fig. 4C-D). All these distributions are highly similar, indicating a structure underlying the interactions among gene/area transcription that is insensitive to both the number and the identity of genes/areas involved in the system.

**Fig. 4.**
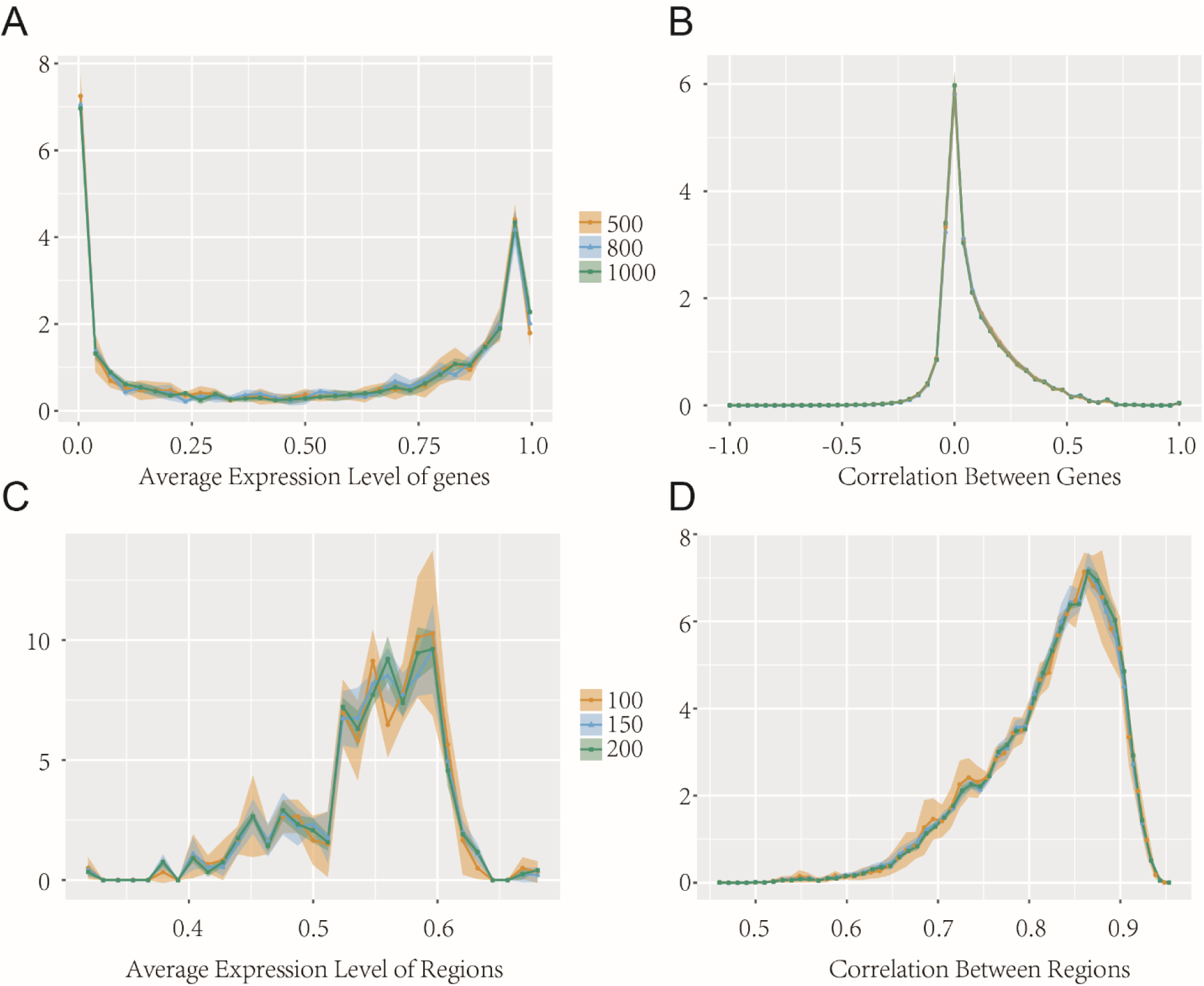
The distribution of the determinants taking control of the pair-wise structures. The distribution of PAA **(A)**, CBA **(B)** with randomly chosen 500, 800, 1000 genes, PAG **(C)**, CGE **(D)** with randomly chosen 100, 150, 200 regions, 5 times each. The numbers of selected genes and regions are color coded. The lines and shaded areas represent the mean ± variance, respectively.

### Coordinated gene expressions shape the architecture of brain networks

Understanding the interaction structures at both the gene and area levels allowed us to explore how coordinated gene expressions could shape the architecture of brain networks. To this end, we ran simulations while changing the expression correlation between genes to examine how these changes would affect the clustering of brain areas in terms of gene expressions. Fig. 5A-C show the clustering results corresponding to a high, intermediate and low level of average gene-gene correlation, respectively. A strong gene-gene correlation tended to form big clusters, while a weak correlation hardly led to any cluster. An intermediate correlation level, which was chosen according to the average gene-gene interactions empirically observed in the AIBS data set, gave rise to complex clustering patterns. To quantify this effect, we adjusted the correlation of genes in a finer resolution and calculated the number of resulting clusters. We found that an average correlation value close to the empirical data led to nearly maximal number of clusters (Fig. 5D). These results were robust to the changes in the number of genes or “areas” used in the simulation (Fig. S7). As the average correlation of genome expression between different areas grew with the expression correlation between different genes (Fig. 5E), our results suggest that an intermediate level of gene-gene correlation leads to a moderate area-area of correlation, which eventually gives rise to a whole brain expression profile allowing the formation of diverse clusters.

**Fig. 5.**
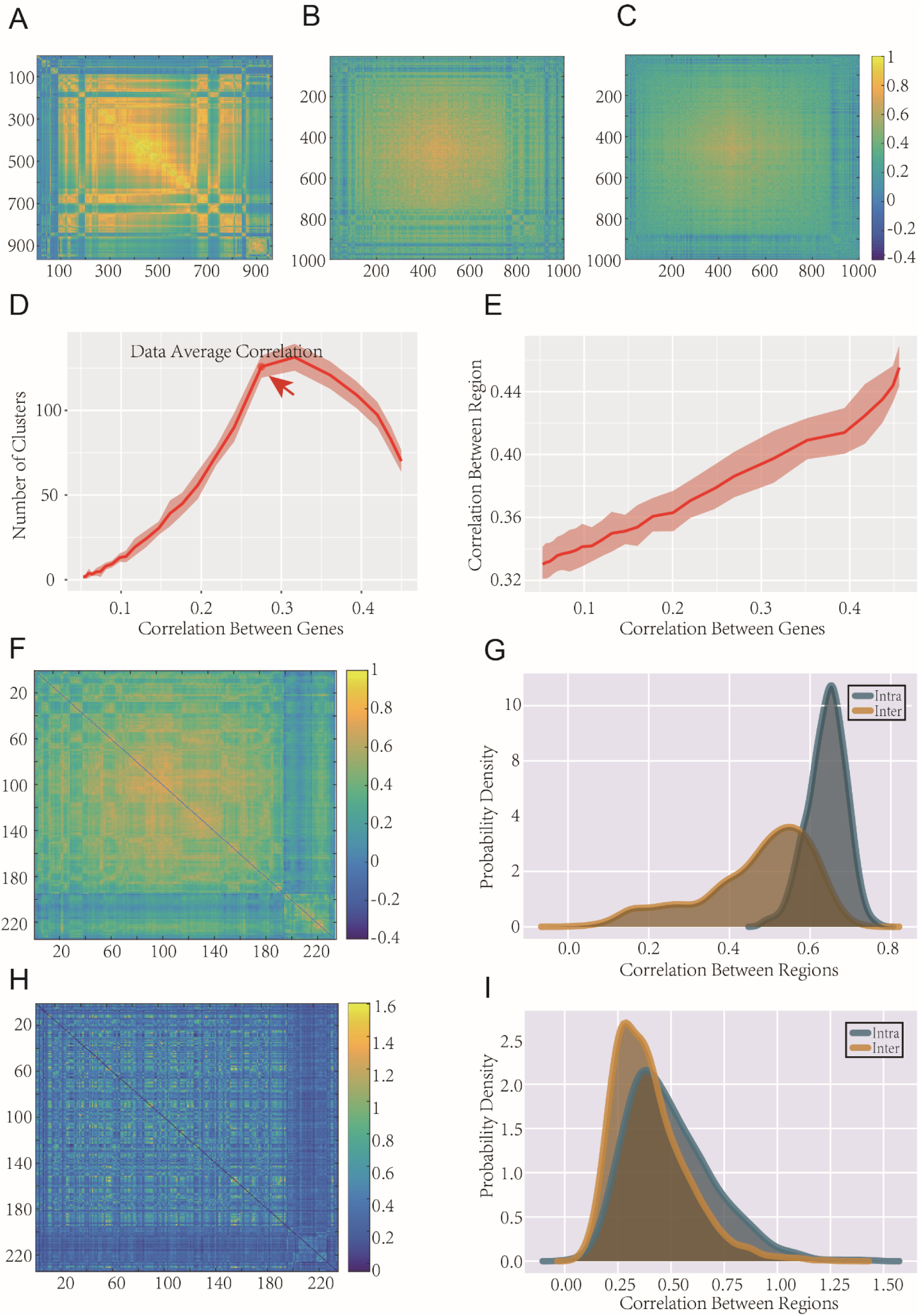
The strength of gene-gene correlations affects the clustering of brain areas. **(A-C)** Correlation matrix showing the clustering results of different regions corresponding to DG model simulations with a high (A), intermediate (B), and low (C) average pair-wise correlation of expression patterns between genes. Color bar, the correlation coefficient of genome expression between areas. (**D**) The number of clusters is plotted as a function of the average correlation of gene-gene correlation for a certain threshold (chosen as 170% of average correlation between areas). Red arrow points at the correlation level corresponding to what was observed in the human transcription data. The red line and shaded area represent the mean ± variance of 10 simulations, respectively. (**E**) The average correlation between areas is plotted as a function of the average correlation between genes. The red line and shaded area represent the mean ± variance of 10 simulations, respectively. (**F**) Correlation matrix showing the clustering results of different regions corresponding to the actual data. **(G)** Distributions of genome expression correlation of areas belonging to the same cluster (intra-cluster, blue) and different clusters (inter-cluster, yellow). (**H**) Correlation matrix showing the functional connectivity of different regions. The order of regions were the same as (F). **(I)** Distributions of functional correlation of areas belonging to the same (intra-cluster, blue) or different transcriptional clusters (inter-cluster, yellow).

Next, we examined the relationship between the clusters of areas with similar genome expression pattern to the brain’s functional connectivity. Specifically, we clustered the brain areas based on their empirically observed expression patterns (Fig. 5F, G) and investigated if these transcriptional clusters are related to areas that are clustered based on correlated neuronal activities [4], [5], [7]. Indeed, we found transcriptional clusters also exhibited clustering property in the functional domain, with the areas within a transcriptional cluster having significantly stronger functional correlation compared to the areas belong to different clusters (Mann–Whitney U test, p<10^−6^) (Fig. 5H, I). Consistent with previous findings [5], [7], these results demonstrate a profound link between the transcriptional and functional relations between brain areas.

Considering that the empirically observed gene-gene correlation gives rise to the maximally diverse transcriptional clusters, our results suggest a possibility that this specific level of gene-gene correlation also contributes to enhancing the brain’s functional richness by facilitating the formation of numerous functional clusters.

## DISCUSSION

In the present study, we provided a quantitative framework to explain the structure of the expression patterns of whole genome for individual areas and the expression profile of the entire brain network. Despite tens of thousands of genes and hundreds of brain areas are involved, we found that at both levels, the pairwise interaction structure can accurately explain the overall expression patterns, suggesting a previously unknown, simple organization that gives rise to well-coordinated gene expressions. Further, based on such a hierarchical structure, we studied how the changes in individual genes’ expressions can affect the properties of the entire brain network through aggregated pair-wise gene-gene, as well as area-area interactions. By numerical simulation, we found that the strength of gene-gene interactions observed empirically allows the nearly maximal number of transcriptionally similar clusters of areas to form, which may account for the functional and structural richness of the brain.

Some methodological limitations might be noteworthy. Firstly, our work was not on specific gene modules or functional brain sub-networks. Although the results provided a basic organization manifesting at the level of the genome and whole brain network, it will be informative for future studies to explore how this structure may vary across individual gene modules or functional brain sub-networks. Secondly, in the data set we analyzed, it didn’t distinct cell types, instead the expression level of many sampled cells in the same site were averaged. We then further average the data across sites that were belonging to the same brain area. Although we demonstrated the latter process did not affect the conclusion (*cf.* fig. S2-6), whether the same principle can be found at the level of single cells awaits future studies to investigate higher resolution data. Finally, the DG model we used works well with binarized data, which has arguably better signal-to-noise ratio than the original, continuous data. However, if one is interested in studying the interaction structure without any binarization process, different parametric model other than the DG model would be required.

Gene expression patterns play a vital role in determining the phenotypes and the behaviors of an organism. A previous study found that expression patterns between genes, rather than aggregated effects of individual genes, determined the phenotypes of the budding yeast [25]. Regarding the brain, at the gene level, it was found that distinct gene expression patterns might account for the different phenotypes of mouse brain and affect animals’ response to seizure [2]. At the level of brain areas, co-expression patterns between areas can reflect and potentially regulate the functional and anatomical structure of the brains in both humans and mice [3]–[5]. Though expression patterns between genes or areas provide vital insights in understanding how the structure and function of brain are shaped by genetic information, the study of expression pattern faces a challenging problem—the curse of dimensionality. That is, even if we only consider a binary state of each gene, for a *N*-gene system, there are in total of 2^*N*^ possible patterns. Here, by using a well-established, data-driven method in studying interaction structure in neural networks [29]–[31], we found that the pair-wise interactions dominant the gene-gene or area-area interactions and largely determine the expression patterns. This means that we do not need to take any high-order complex interactions among genes or areas of the brain into account, instead we only needed to consider the pair-wise interactions and expression probability of genes or areas. As a result, it dramatically reduces the complexity of the transcription patterns of genes and areas, i.e, from 2^*N*^ to *N*^2^, thereby opening the possibility to comprehensively study the relationship between transcription patterns and the structure as well as the function of the brain.

Also, these results have important methodological implications as the pair-wise approach is widely used in analyzing transcriptional data. At the gene level, previous studies identified biologically relevant modules across the brains with the weighted gene co-expression network analysis (WGCNA) [38], [39] by grouping genes with strong pair-wise covarying patterns into modules [2], [4], [21]. At the area level, it is also common to use pair-wise correlation between areas or tissues to construct the cortical-cortical co-expression network[6], [7]. For a system with strong high order interactions (HOIs), it has been demonstrated that the pair-wise approaches will lead to incorrect conclusion regarding the true interaction structure [40], thus the absence of strong HOIs discovered here provides a solid foundation on which previous studies regarding gene expressions can stand.

It is noteworthy to clarify the relation between the HOIs absent in the current study to those usually reported in other graph-theoretical studies of interactions. Higher order interactions are commonly found in studies of complex networks in fields ranging from social sciences to biology [41]–[44]. These structures, usually known as motifs, can be understood as aggregated pair-wise interactions. It has been found that such interactions may have important roles in mediating or even determining the global behaviors of the systems [45], [46]. In contrast, the HOIs we studied here refer to the ones that cannot be reduced to any pair-wise interactions [e.g., the Exclusive OR or XOR function [47]]. They have been studied in different scales of neural networks in various species [29], [48], [49] as well as in group activities of rodents [50]. Thus, the current conclusion of the absence of HOIs is not contradictory to previous findings of motifs in gene-gene interactions [46], [51].

The human brain is a complex modular structure, with numerous structurally and functionally different areas (modules) orchestrated to support diverse functions. Such a modular structure is fundamentally controlled by gene expressions. A previous study found that even though transcriptional regulation was varied enormously according to the anatomical location, different regions and their constituent cell types displayed highly conserved molecular signatures between individuals, suggesting the conservative transcriptional regulation [2]. It has also been found that gene expressions are correlated with functional connectivity, suggesting a strong link between gene expressions and the functions of individual brain areas [4]–[7]. These previous findings shed light on interpreting our result that the empirically observed level of gene-gene correlation gives rise to the nearly maximally diverse transcriptional clusters of areas. That is, such diverse transcriptional clusters may lead to numerous brain areas, which collectively form the complex modular architecture of the brain and ensure its functional richness.

In the simulation with differrent levels of gene-gene correlations, there exists a two-stage process organized hierarchically: 1). changing the interactions between genes resulted in altered genome expression patterns for individual areas, which further changed the expression interactions between areas; 2). different interactions between areas ultimately led to re-organization of the entire brain network in terms of gene expressions, determining e.g., how many clutters can be formed. Thus, by elucidating the interaction structure among gene expressions, we provide a framework that link activities of individual genes at the microscopic level all the way to the macroscopic properties of the entire brain, which may be instrumental to understand how genetic information modulates the structure and function of the brain. It may also assist the studies on both the pathogenesis of some complex brain disorders and the mechanisms underlying specific brain functions. For example, through comparing the expression data between patients of neurodegenerative diseases to the healthy controls, many differentially expressed genes (DEG) have been documented [52]–[54]. Also, when it comes to specific functions of the brain, such as synaptic functions [2], regulating dopamine signals subserving the working memory [55], etc. lots of genes relate to these specific functions have been detected by Gene Ontology (GO) or other analyses [56], [57]. These studies illustrated a fairly complicated pictures regarding the potential mechanism of brain disease and function. However, according to the present framework, the changes in minority of key genes may propagate through pair-wise interactions to affect a large number of genes and gives rise to the apparent complexity that many genes are involved. Indeed, hub genes or specific modules were found to be related to some vital biological function and disease. For example, a precious study [58] found that a noncoding RNA DGCR5, identified as a hub gene, played a potential role in regulating certain Schizophrenia related genes. TREM2, another hub gene found previously [59], mediated changes in the microglial cytoskeleton necessary for both phagocytosis and migration. Our finding might provide a parsimonious explanation that minority of hub or key genes’ variation could eventually lead to alterations in the whole system’s behavior or function. In that view, seemingly complicated mechanisms of brain diseases and functions might be triggered by a few key genes, which may enlighten future studies to trace these critical genes. Furthermore, some brain disorders sharing symptoms and substantial epidemiological comorbidity have some common genetic variations, such as schizophrenia and bipolar disorder [60], anxiety disorder and major depression [61], etc. It has been found the root of these diseases have significant similarity at the molecule level [62]. Our work suggests a possibility that these diseases might be intrigued by similar key genes, which give rise to different brain diseases by differential regulation of the pair-wise gene-gene interactions.

We also found that the distribution of the determinants, the PAA, CBA, PAG, CGE, of the structure at the two levels were ultimately converged with the increase of the number of the genes or areas and not affected by specific genes or areas, suggesting the determinants were also shaped by some underlying constrains, which is beyond of our reach. We believe these fixed distribution may provide a baseline or template to distinguish diseases, which is also the aim of [4]. We hope these findings might offer some inspiration when it refers to the relationships between structure, function, development, evolution and disease.

## Author contributions

JH and SY conceived the study, JH performed the analyses, ZY and TJ provided tools for analyses, JH and SY wrote the manuscript.

## Data and materials availability

All data, code, and materials used in the analysis are available upon request for purposes of reproducing or extending the analysis.

