## Supplementary material for "Pair-wise Interactions in Gene Expression Determine a Hierarchical Transcription Profile of the Human Brain": methods

**This PDF file includes:**

Materials and Methods

Figs. S1 to S7

Captions for Data S1

### Materials and Methods

1. **Data**

### 1.1 Human brain transcription data set

The human brain microarray data provided by the Allen Institute [1], [2] were analyzed in the present study. The data set was obtained from 6 subjects (5 males and 1 females; 24-57 years old), including two whole brains and 4 left hemispheres. In total, there are 3702 different sites were sampled, which were distributed across the whole brain. For each site, there were 58692 probes used to estimate the expression level of all genes. This data set contains both the raw data representing continuous expression levels and their binarized counterparts representing the active (1) and inactive (0) state of individual genes. The complete datasets can be downloaded from <http://human.brain-map.org/static/download>, and relevant documentations can be found at <http://help.brain-map.org/display/humanbrain/Documentation>.

From the total of 58692 probes, we chose the probes uniquely corresponding to 16906 genes according to a previous study [3]. The complete list of the probes can be found in the supplementary data file of [3].

### 1.2 Structural and functional connectivity of the human brain

The Human Brainnetome Atlas (HBA), released by the Brainnetome Center, with 210 cortical and 36 subcortical regions, provides a detailed parcellation of the human brain and the connection information among different brain areas [4]. The HBA was constructed from the data released by the Human Connectome Project (HCP). Based on HBA, the functional connectivity was obtained by calculating the Pearson’s correlation coefficient between the time series of BOLD signals during the resting state of each region. Both the structural and functional connectivity data can be downloaded from <http://atlas.brainnetome.org/download.html> (See [4] for more details).

### 2． Data preprocessing

### 2.1 Registering locations of individual samples to different regions

The coordinates in the MNI space of each sample in the transcription data used here were converted into the voxel space of the HBA, using the information of the voxel size and the spatial origin coordinates of the HBA in the MNI space. 1578 samples were discarded from further analyses as they were not located within the 246 areas covered by HBA (cerebral cortex and subcortical nuclei). As a result, a total of 2124 sample sites remaining were then mapped into different regions in HBA. In all, 12 areas included in the HBA have not registered any samples, and thus were not included, resulting in a total of 234 regions in the present analyses. The median number of samples for a given region was 7 (ranging from 1 to 75) (Fig. S1 for the distribution of the sample numbers in a given region).

### 2.2 Obtaining average expression profile for individual regions

The expression patterns of individual areas were determined according to the majority of samples that registered to the specific areas. That is, if ≥ 50% of samples exhibited active expression of a specific gene in the same area, this gene was considered to be active for this area.

The results shown in the main text were based on aggregated data from all six subjects. Highly similar results were obtained from individual subjects (Fig. S2-5). In addition, highly similar results were obtained if the analyses were carried out at individual sample sites, i.e., without merging the samples belonging to the same area (Fig. S6).

### 3. Parametric models used

### 3.1 The independent model (IM)

The independent model only uses the probability of each gene to be active or the proportion of active genes in each brain area as its input for prediction. In other words, the model assumes expression patterns of different genes or brain areas are independent with each other. Taking two genes for example, if gene A, with the probability of $P_{A}$ to be active, is independent of gene B, with the probability of $P_{B}$to be active, then the probability of gene A and B simultaneously to be active $P_{AB}$is:

$P_{AB}= P_{A}\times P_{B}$	(1)

The prediction of the IM was obtained by running 10000 times of Bernoulli experiments with each gene to be active or the proportion of active genes in each brain area set to be the same as empirically observed.

### 3.2 Dichotomized Gaussian (DG) Model

The DG model [5], [6] uses the same information as the IM plus the expression covariance between two genes or areas as its inputs for prediction. The process that the DG model generates samples can be divided into two steps: 1) given the number of genes or areas as *N*, we drew 10000 samples from the *N*-dimensional Gaussian distribution, with the mean value of $\gamma$ and the covariance of $\Lambda$, 2) for each sampled *N*-dimensional vector, we binarized it with the threshold of 0, resulting in the binarized samples with the mean of $r$ and pairwise covariance of $\Sigma$. $r$ and $\Sigma$ were set to be the same as empirically observed. To achieve this, $\gamma$ and$\Lambda$ were obtained by numerically solving the following equations:

$r_{i}= \Phi\left( \gamma_{i} \right)$	(2)

$\Sigma_{ii}= \Phi\left( \gamma_{i} \right)* \Phi\left( {-\gamma}_{i} \right)$	(3)

$\Sigma_{ij}= \Phi_{2}\left( \gamma_{i},\gamma_{j},\Lambda_{ij} \right)-\Phi\left( \gamma_{i} \right)*\Phi\left( \gamma_{j} \right)$	(4)

where *i* and *j* represent different genes/areas, $\Phi$ represents the cumulative distribution of a univariate Gaussian with the mean of 0 and the variance of 1, and $\Phi_{2}$ represents the bivariate counterpart of $\Phi$with the covariance of $\Lambda_{ij}$.

### 4. Hierarchical clustering

Hierarchical clustering [7] built a nested tree of clusters based on the distance between different nodes. In the hierarchical clustering tree, the root node has the maximal cluster, which includes all the leaf nodes, while the leaf node has the minimal cluster, which contains merely itself. Each child node has the same class label as its parent node. In the present study, we took each brain area (or each sample in the DG model simulation at the gene level) as a node and used the bottom-up merging method to construct a hierarchical clustering tree. Specifically, we carried out the following steps: 1) calculate the distance matrix between nodes as (1-*C*)/2, where *C* represents the correlation matrix; 2) choose the nearest two nodes from the node set; 3) merge them to create a new node; 4) add the new generated node into the node set and delete the two old nodes; 5) repeat the steps from 1 to 4 until only one node remains, which would be the root node. The Matlab function *linkage* was used for hierarchical clustering and *optimalleaforder* was used for optimizing the clustering[8].

In the results, a cluster is defined as the group of areas with the average correlation *r* higher than a predefined threshold value *t*, which is in turn measured by the average of the entire correlation matrix *R* (e.g., *t*= 1.5*R*, 1.6*R* and 1.7*R*). Specifically, we used a two-step process to define the clusters, i.e. the searching sub-process, and the merging sub-process. In addition, the group of areas had three status, i.e. non-cluster (the areas of it cannot be considered as a cluster), possible cluster (the cluster could be merged with an adjacent cluster to form a bigger possible cluster), and definite cluster (the cluster could not be merged with any other adjacent possible cluster). In the searching sub-process, we started from a start area with a possible cluster size *s*=16 to detect a possible cluster (*r* ≥ *t*). If *r* < *t*, *s* would be reduced to its half value and update the possible cluster size. This process repeated until 1) *s* < 2, in this case the remaining area was considered as a non-cluster. The search program for a possible cluster re-ran and the next area was taken as the new start area. The possible clusters searched before would be taken as the definite cluster. 2) *r* ≥ *t* and *s* ≥ 2, in this case the corresponding areas were considered to belong to a possible cluster, and the search moved to the next area for another possible cluster. In the merging sub-process, we merged two adjacent possible clusters into a bigger possible cluster if the bigger one had *r* ≥ *t*, otherwise the former cluster was taken as a definite cluster. The process repeated the searching and merging until the last brain area was processed.

### 5. Influence of gene correlation on both area correlation and cluster formation

To understand how coordinated gene expressions could ultimately influence the overall features of the entire brain network, we changed the expression correlation between genes to examine the clustering results of brain areas. The procedure was the following: 1) randomly chose 400 genes to run the DG model at the gene level to calculate the corresponding $\gamma$ and $\Lambda$; 2) gradually increased or reduced $\Lambda$ to get a new $\Lambda'$; 3) calculated the nearest positive define matrix [matrix](javascript:;) $\Lambda''$ of $\Lambda'$ according to [9]; 4) re-ran the DG model simulation, with $\gamma$ and $\Lambda''$ to generate 1000 samples (representing 1000 brain areas) 5) calculated the average correlation between samples; 6) hierarchically clustered the new samples according to the correlation between these samples; 7) obtained cluster numbers based on different thresholds (150%, 160% and 170% of the actual average correlation between areas). In addition, we deleted the clusters with small *s*, i.e. clusters including only a few areas. Specifically, we repeated steps 5-7 after generating 1000 samples by the IM model simulation and chose the maximum number of areas in these clusters as the threshold *s* to delete all clusters smaller than it from the simulated data.

### 6. Performance measurements of models

### 6.1 JS divergence

Here, we used JS divergence to quantify the distance between two distributions obtained from actual data and models’ prediction. JS divergence is built on the Kullback–Leibler (KL) divergence, which is another asymmetric distance measurement of distributions. Generally, JS divergency is symmetric and its value ranges from 0 to 1. Canonically, P represents the true distribution of the data, while Q represents its theoretical distribution, in other words, model distribution or approximate distribution. In this regard, the JS divergency between P and Q can be defined as:

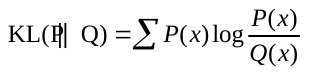
,

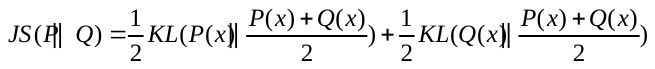
 ,

where
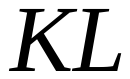
represents the KL divergence.

### 6.2 RED

Here, we used the RED to quantify the percentage of the characteristics that can be captured by DG model. Generally, for an arbitrary network of
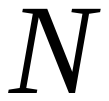
elements, if possibility, there exists
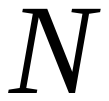
 kinds of correlations, like pair-wise, triplet or other more complex ones. As a result, we construct a distribution of maximum entropy of the network
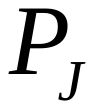
, measure how much amount can be captured by up to J-order correlations. Specifically,
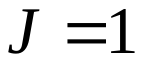
 means all the genes are independent and
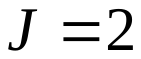
 means the network can be captured by up to pair-wise interactions between genes. However, if
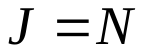
, the network will allow all arbitrarily complex interactions which can’t be numerated with the increase of
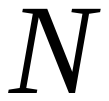
. Here, the entropies
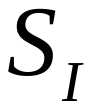
 decrease monotonically toward the true entropy S:
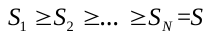
.
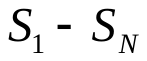
 measures the summary amount of information which can be captured by all kinds of correlations. In addition, the information contribution of J-order interactions can be captured by
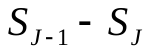
. As a result, the contribution of up to J-order correlations can be measured by
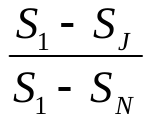
 , which is named as RED here.

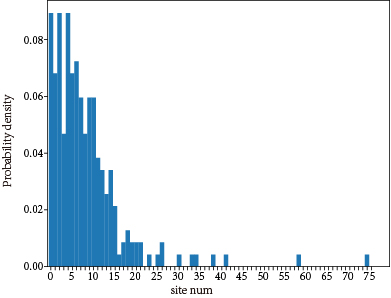

**Figure S1. Distribution of the number of sites assigned to a given region of HBA.**

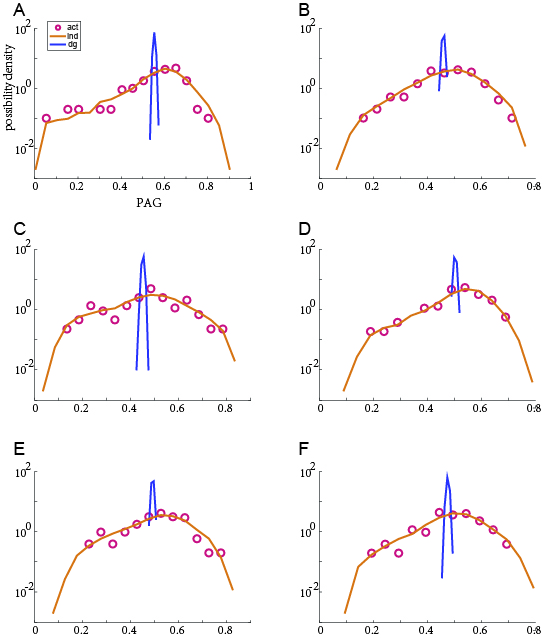

**Figure S2. PAG of individual subjects.** (**A-F**) The distribution of PAG of each of the six subjects (red circles) and the corresponding predictions based on the IM (blue) and DG (yellow) model.

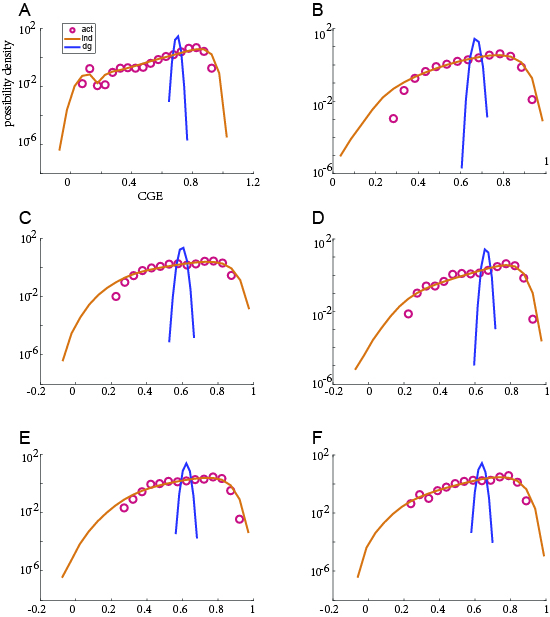

**Figure S3. CGE of individual subjects.** (**A-F**) The distribution of CGE of each of the six subjects (red circles) and corresponding predictions based on the IM (blue) and DG (yellow) model.

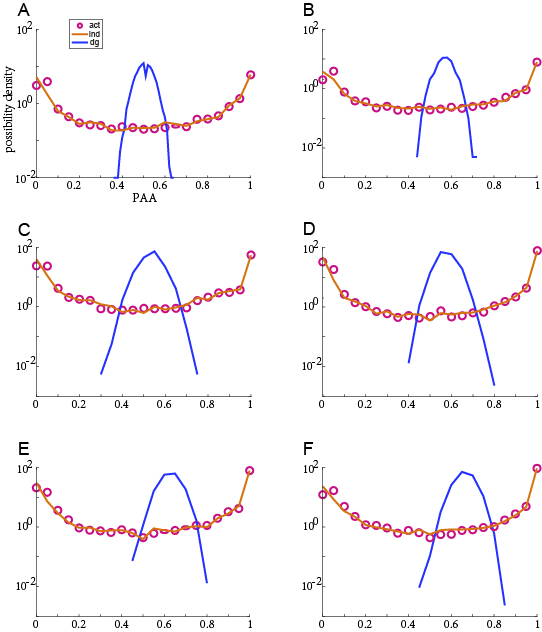

**Figure S4. PAA of individual subjects.** (**A-F**) The distribution of PAA of each of the six subjects (red circles) and corresponding predictions based on the IM (blue) and DG (yellow) model.

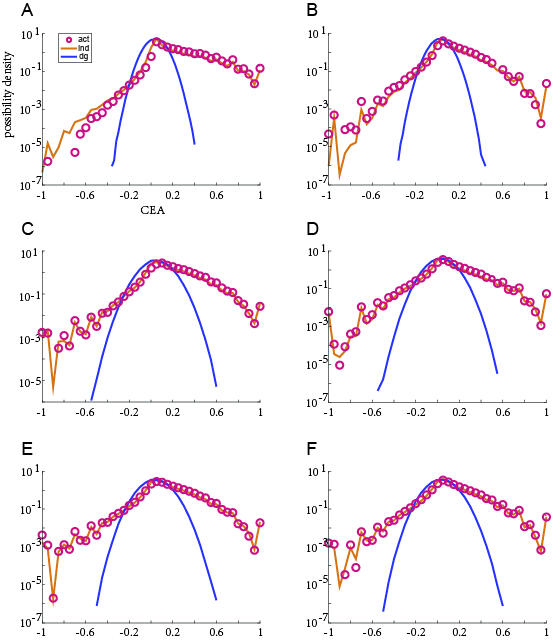

**Figure S5. CEA of individual subjects.** (**A-F**) The distribution of CEA of each of the six subjects (red circles) and corresponding predictions based on the IM (blue) and DG (yellow) model.

.

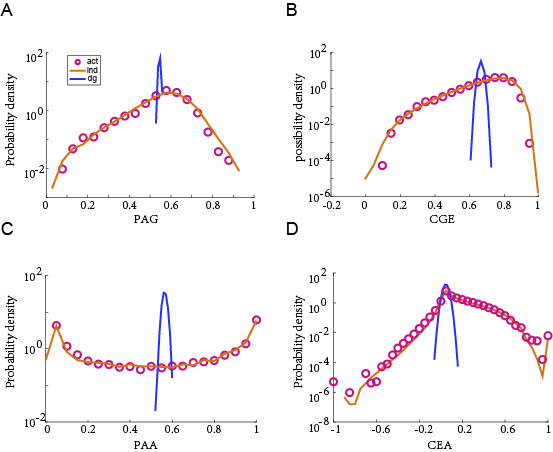

**Figure S6. Evaluation of different models at the level of individual sample sites.** Based on aggregated individual samples, i.e., 2124 sites in the brain, the distribution of PAG (**A**), CGE (**B**), PAA (**C**), CEA (**D**) (red circles) and the corresponding predictions based on the IM (blue) and DG (yellow) model.

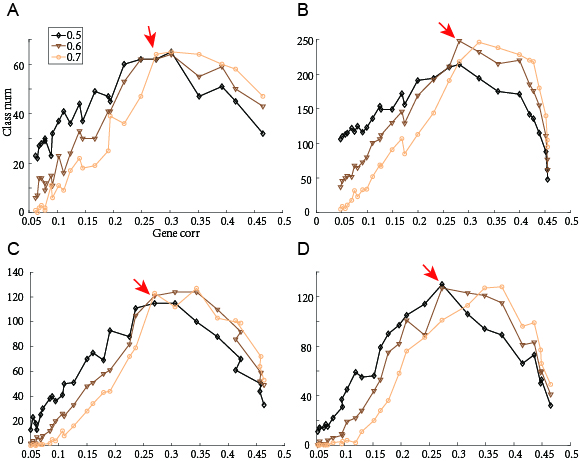

**Figure S7. Influence of gene correlation on area correlation and cluster formation with different numbers of genes or samples.** (**A-B**) the curves of a randomly selected 400-gene system with 500 and 2000 samples, respectively. (**C-D**) the curves of randomly selected 600- and 800-genes systems with 1000 samples, respectively (different colors represent the threshold of 150%, 160%, 170% of average correlation between areas, respectively).

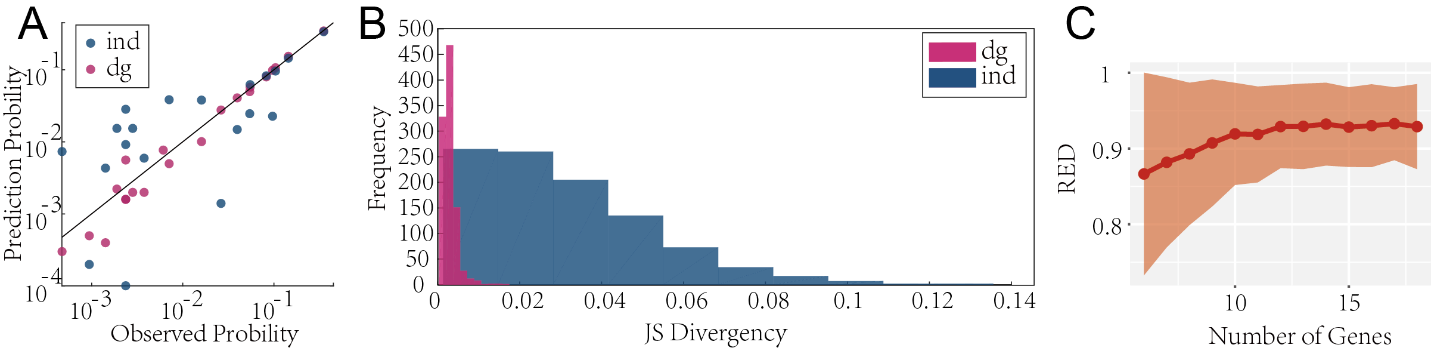

**Figure S8. The expression pattern of randomly chosen 6 genes with 2124 samples site can be accurately predicted by pair-wise interactions between genes. (A)** using a random chosen 6 genes with 2124 samples, the possibility of each expression pattern observed is plotted against that predicted from the DG model (red dots) and IM (blue dots). Black line shows equality. **(B)** Distributions of JS divergences between the possibility of expression patterns observed and that predicted by DG model (red) and IM (blue); data from 1000 groups of randomly chosen 6 genes with 2124 samples. **(C)** Randomly chose 6~18 genes with 2124 samples, 1000 times each. Number of genes are plotted against REDs. The line and shaded areas represent the mean ± variance, respectively.
